# Gene Drive Mosquitoes Can Aid Malaria Elimination by Retarding Plasmodium Sporogonic Development

**DOI:** 10.1101/2022.02.15.480588

**Authors:** Astrid Hoermann, Tibebu Habtewold, Prashanth Selvaraj, Giuseppe Del Corsano, Paolo Capriotti, Maria G. Inghilterra, Temesgen M. Kebede, George K. Christophides, Nikolai Windbichler

## Abstract

Gene drives hold promise for the genetic control of malaria vectors. The development of vector population modification strategies hinges on the availability of effector mechanisms impeding parasite development in transgenic mosquitoes. We augmented a midgut gene of the malaria mosquito *Anopheles gambiae* to secrete two exogenous antimicrobial peptides, Magainin 2 and Melittin. This small genetic modification, capable of efficient non-autonomous gene drive, hampers oocyst development in both *Plasmodium falciparum* and *Plasmodium berghei*. It delays the release of infectious sporozoites while it simultaneously reduces the lifespan of homozygous female transgenic mosquitoes. Modeling the spread of this modification using a large-scale agent-based model of malaria epidemiology reveals that it can break the cycle of disease transmission across a range of endemic settings.

**One sentence summary:** We developed a gene drive effector that retards *Plasmodium* development in transgenic *Anopheles gambiae* mosquitoes via the expression of antimicrobial peptides in the midgut and which is predicted to eliminate malaria under a range of transmission scenarios.

## Main text

Malaria remains one of the most devastating human diseases. A surge in insecticide-resistant mosquitoes and drug-resistant parasites has brought a decades-long period of progress in reducing cases and deaths to a standstill (*1*). Despite the availability of the first WHO approved malaria vaccine (*2*) the necessity to develop alternative intervention strategies remains pressing, particularly if malaria elimination is to remain the goal. Gene drive, based on the super-Mendelian spread of endonuclease genes, is a promising new control strategy that has been under development for over a decade (*3*). Suppressing mosquito populations by targeting female fertility has remained a prime application of gene drives and to date specific gene drives have been shown to eliminate caged mosquito populations (*4–6*). Gene drives for population replacement (or modification), designed to propagate antimalarial effector traits, have also seen significant development in past years (*7–9*). To date, a range of antimalarial effectors and tissue-specific drivers have been tested in transgenic mosquitoes, and some of them have been shown to reduce *Plasmodium* infection prevalence or infection intensity (*10–23*). However, the pursuit of novel and effective mechanisms is ongoing, especially in *A. gambiae* where effectors so far have shown only moderate reductions in parasite transmission (*12, 15, 20–22*).

Antimicrobial peptides (AMPs) from reptiles, plants or insects have long been considered putative antimalarial effectors and have been tested *in vitro* and *in vivo* for their efficacy against different parasite life stages (for reviews see (*24–27*); for more recent studies see (*28, 29*)). Although AMPs are very diverse in sequence and structure, many are cationic and amphiphilic and thus tend to adhere to negatively charged microbial membranes and to a much lesser extent to membranes of animal cells (*30*). Permeabilization mechanisms have been proposed, which rely on either pore formation or accumulation of peptides on the microbial surface causing disruption in a detergent-like manner (*31*). A subset of AMPs has been suggested to act by mitochondrial uncoupling, interfering directly with mitochondria-dependent ATP synthesis (*32–34*). Two such peptides, Magainin 2, found within skin secretions of the African claw frog *Xenopus laevis*, and Melittin, a primary toxin component of the European honeybee *Apis mellifera*, have been shown to both form pores on the microbial membrane (*35, 36*) and trigger uncoupling of mitochondrial respiration (*37–40*). Intrathoracic injection of Magainin 2 into *Anopheles* mosquitoes has been demonstrated to cause *Plasmodium* oocyst degeneration and shrinkage and a consequent reduction in the number of sporozoites released (*41*), while a transmission blocking effect of Magainin 2 has also been revealed when spiked into gametocytaemic blood at a 50 μM concentration (*29*). Similarly, Melittin has been shown to reduce the number and prevalence of *P. falciparum* oocysts in spike-in experiments at concentrations as low as 4 μM (*28, 29*), while expression of Melittin in transgenic *A. stephensi* mosquitoes as a part of a multi-effector transgene, additionally including the AMPs TP10, EPIP, Shiva1 and Scorpine, has led to a significant reduction in oocyst prevalence and infection intensity (*23*).

Here, we augmented two host genes of *A. gambiae* to co-express Magainin 2 and Melittin following the previously described integral gene drive (IGD) paradigm (*42*). This allowed for minimal genetic modifications, capable of non-autonomous gene drive, to be introduced into the host gene loci making full use of the gene regulatory regions for controlling tissue-specific expression of the AMPs. We utilized the previously evaluated zinc carboxypeptidase A1 (CP, AGAP009593) and the AMP Gambicin 1 (Gam1, AGAP008645) as host genes for the exogenous AMP integration (**Fig. 1A**). The transcriptional profile of these genes was expected to drive expression of the AMPs in the mosquito midgut upon ingestion of a bloodmeal (CP) or in the anterior gut (Gam1). The use of 2A ribosome-skipping peptides as well as secretion signals guaranteed separate secretion of the exogenous AMPs and host gene products. We also replaced the signal peptides and prepropeptides of Magainin 2 and Melittin with the endogenous secretion signals of *A. gambiae* Cecropin 1 and 2 genes, respectively. An intron harboring the guide RNA-module that enables non-autonomous gene drive and the fluorescent marker-module required for transgenesis was introduced within the Melittin coding sequence. Transgenesis of *A. gambiae* G3 strain via CRISPR/Cas9-mediated homology-directed repair (HDR) and subsequent removal of the GFP transformation maker resulting in the establishment of homozygous markerless strains, designated as Gam1-MM and MM-CP, were performed as previously described (*42*) (**Fig. 1A**). We validated and tracked the correct transgene insertion by genomic PCR (**Fig. 1B**) and confirmed the expression of both CP and Gam1 host genes and the inserted AMP cassette by RT-PCR (**Fig. 1C**). Sequencing of the cDNA amplicon over the splice site of the artificial intron revealed the expected splicing pattern in 89.9% of all MM-CP reads but only in 65.3% of all Gam1-MM reads (**Fig. 1D**). A cryptic splice site resulting in a loss of additional 25bp from the Melittin coding sequence accounted for most of the unexpected splicing events (**Fig. S1**).

**Figure 1.**
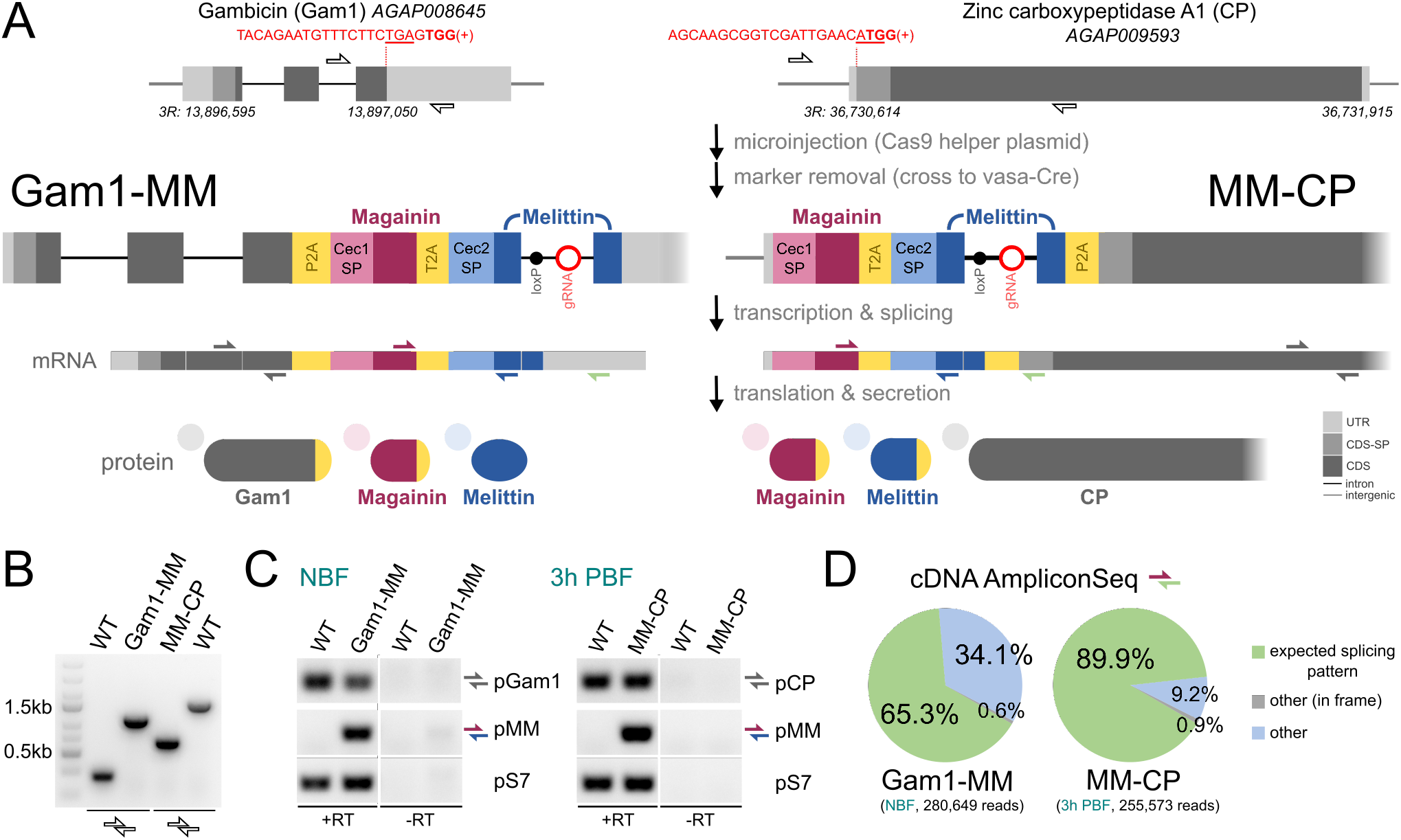
Generation of gene drive effector strains expressing AMPs. **(A)** Schematic showing the design and integration strategy of the effector cassette coding for Magainin 2 and Melittin at the endogenous loci Gam1 and CP. The gRNA target sequences (red), or gRNA module (red circle) are indicated including the protospacer adjacent motif (bold) and the Stop and Start codons (underlines). Coding sequences (CDS) and signal peptides (CDS-SP) are indicated by light shading. Half arrows indicate primer binding sites for genomic PCR and RT-PCR. **(B)** PCR on genomic DNA of 15 pooled homozygous Gam1-MM, MM-CP or wild type individuals. **(C)** RT-PCR of midguts from wildtype (WT), Gam1-MM and MM-CP mosquitoes that were either non-blood-fed (NBF) or dissected three hours post blood-feeding (3h PBF). **(D)** Analysis of cDNA amplicons over the splice site subjected to next generation sequencing showing the predicted splicing outcomes for strains Gam1-MM and MM-CP.

Next, we performed infection experiments with the *P. falciparum* NF54 strain to determine the effect of these modifications on parasite transmission (**Fig. 2A**). Both transgenic strains showed a significant reduction in the midgut oocyst loads on day 7 post infection (pi; **Fig. 2B**). While only few oocysts of the MM-CP strain had the expected size, a closer examination of infected midguts revealed the presence of many smaller structures possibly representing stunted or aborted oocysts (**Fig. 2C**), prompting a more detailed investigation of this strain. We performed infections and quantified the number and diameter of all oocyst-like structures on days 7, 9 and 15 pi. Given that nutritional stress is a factor that recently emerged as causing stunting of oocysts (*43, 44*), a group of mosquitoes were provided a supplemental bloodmeal on day 4 pi. We found that the small oocyst-like structures were indeed stunted oocysts that grew over time, and that the supplemental bloodmeal further boosted their growth (**Fig. 2D**). Overall oocysts infecting the MM-CP strain were significantly smaller than in the wildtype control by an average of 47.8%, 41.1% and 59.8% on days 7, 9 and 15 pi, respectively (**Fig. 2D**). This was also the case for the cohort that received an additional bloodmeal, but the difference with wildtype controls decreased over time to 50.4%, 24.9% and 18.6% on days 7, 9 and 15 pi, respectively. We repeated this experiment with the rodent parasite *P. berghei* to determine if the observed infection phenotype would also occur with a different parasite species and under different environmental conditions. Using a GFP-labeled *P. berghei ANKA* 2.34 strain, fluorescent imaging revealed clearly stunted oocysts that on day 14 pi were about two-times smaller than in the control **(Fig. 2E, F)**.

**Figure 2.**
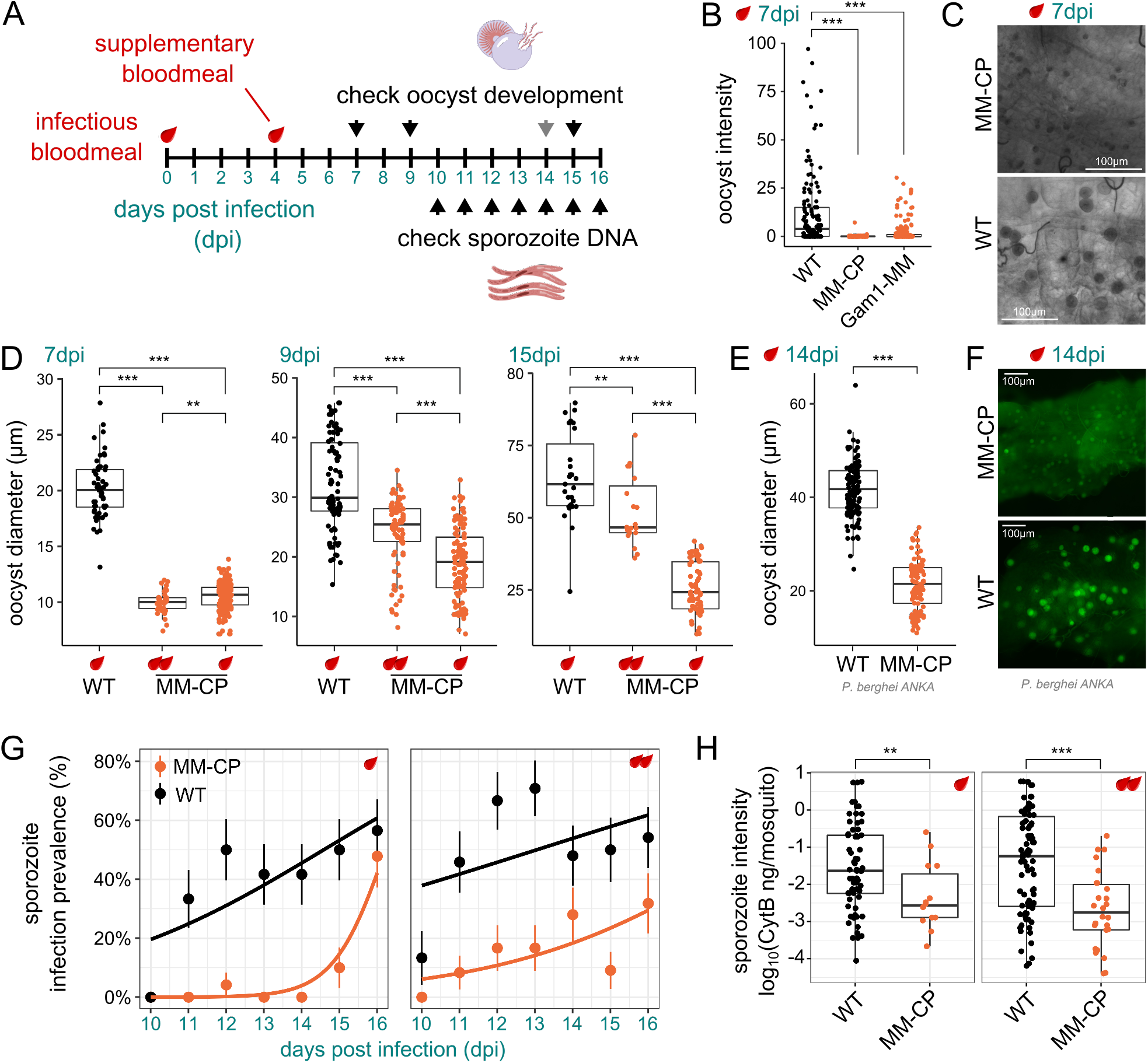
*Plasmodium* infection experiments. **(A)** Schematic overview of *Plasmodium* infection experiments. **(B)** *P. falciparum* oocyst intensity 7 days pi (dpi) in midguts from wildtype (WT), MM-CP and Gam1-MM mosquitoes dissected. Data from three biological replicates was pooled and statistical analysis was performed using the Mann-Whitney test. **(C)** Bright-field images of midguts showing typical oocysts in WT and MM-CP mosquitoes at 7 dpi. **(D)** Quantification of *P. falciparum* oocyst diameter in WT and MM-CP mosquitoes 7, 9 and 15 dpi from 3 pooled biological replicates. Note, that many oocysts in WT mosquitoes had ruptured on day 15 pi. Quantification of oocyst diameter **(E)** and fluorescent imaging **(F)** of oocysts in WT and MM-CP mosquitoes infected with *P. berghei* at 14 dpi. Sporozoite prevalence 10 to 16 dpi **(G)** and infection intensity across all days **(H)** was measured by qPCR of the *P. falciparum* Cyt-B gene in dissected heads and thoraces of individual MM-CP and WT mosquitoes (10-16 dpi). Only mosquitoes positive for oocyst DNA in the midgut were included in the analysis performed in two biological replicates. Statistical analysis in panels D, E & H was performed by a t-test assuming unequal variance. Statistical analysis in panel G was performed using a generalized linear model with a quasibinomial error structure where strain (p=8.35e-06), dpi (p=7.15e-4) but not bloodmeal status (p= 0.0894) were found to be significant coefficients. In all panels, the provision of a supplemental bloodmeal 4 dpi is indicated by an additional blood drop. Significance codes *(p≤0.05), **(p≤0.01), ***(p≤0.001) and ns (not significant).

We reasoned that the detected retardation of oocyst development would in turn cause a delayed release of sporozoites into the mosquito haemocoel and subsequent infection of the salivary glands. To determine the sporozoite load over time, we determined the abundance of parasite DNA on days 10 to 16 pi by quantifying the *P. falciparum* cytochrome B (Cyt-B) gene in the head and thorax of single mosquitos via qPCR, considering only mosquitoes that were positive for parasite DNA in the midgut. We found that sporozoite infection prevalence, *i.e.*, the rate of head and thorax samples with amplification above the amplification cycle threshold, was significantly reduced in MM-CP mosquitoes (**Fig. 2G**) when compared to wildtype (on average by 77.9% across timepoints) with a significant number of positives detectable only on day 16. Although a supplemental bloodmeal accelerated sporozoite release in the MM-CP strain by 4-5 days (now detected on days 11-12 pi), overall sporozoite prevalence was still reduced significantly relative to the control by 67.8% on average. This suggested that sporozoite release in the MM-CP strain was delayed under both nutritional conditions. We also analyzed the relative parasite DNA content in positive samples as a proxy for sporozoite numbers and found that these were significantly higher in wildtype compared to MM-CP mosquitoes both after a single (14.0-fold) or double (37.8-fold) bloodmeals **(Fig 2H)**. To rule out any founder effect in the homozygous MM-CP strain, we outcrossed MM-CP mosquitoes to mosquitoes of the *A. gambiae* Ifakara strain. After an F1 sibling cross, F2 mosquitoes were provided an infected bloodmeal and dissected on day 9 to assess the midgut oocyst size and be genotyped by PCR for the presence of the transgene. The results showed that the effect on parasite development was indeed attributable to the presence of the transgene and that the developmental retardation of oocysts was reduced in hemizygous individuals (**Fig. S2**).

Next, we measured fitness parameters of MM-CP mosquitoes. The results showed a 14.1% difference in the number of eggs laid (fecundity; p=0.0299, two-sample t-test assuming unequal variances) but unaffected larval hatching rates (fertility) between the MM-CP and wildtype control mosquitoes (**Fig. 3A, B**). Pupal sex ratio (**Fig. 3C**) and pupation time did not significantly deviate between MM-CP and control mosquitoes (**Fig. S3**). However, a significant effect on the lifespan of sugar-fed mosquitoes was detected for MM-CP females (median lifespan 15 days) and to a lesser extent in males (23 days) compared to control females (26 days) and males (27 days; **Fig. 3D**). We repeated this experiment with females now also been provided regular bloodmeals. As CP is only weakly expressed in the sugar-fed female midgut but strongly induced following a blood feed, any effect of the transgene was expected to be elevated by the bloodmeals. As before, to rule out that inbreeding accounted for this effect, we first performed a backcross to the Ifakara strain and genotyped individual F2 mosquitoes at the end of the experiment. The results confirmed a significant lifespan reduction in homozygous MM-CP females under these conditions, but no significant effect was detected in hemizygous individuals compared to non-transgenic controls (**Fig. 3E**).

**Figure 3.**
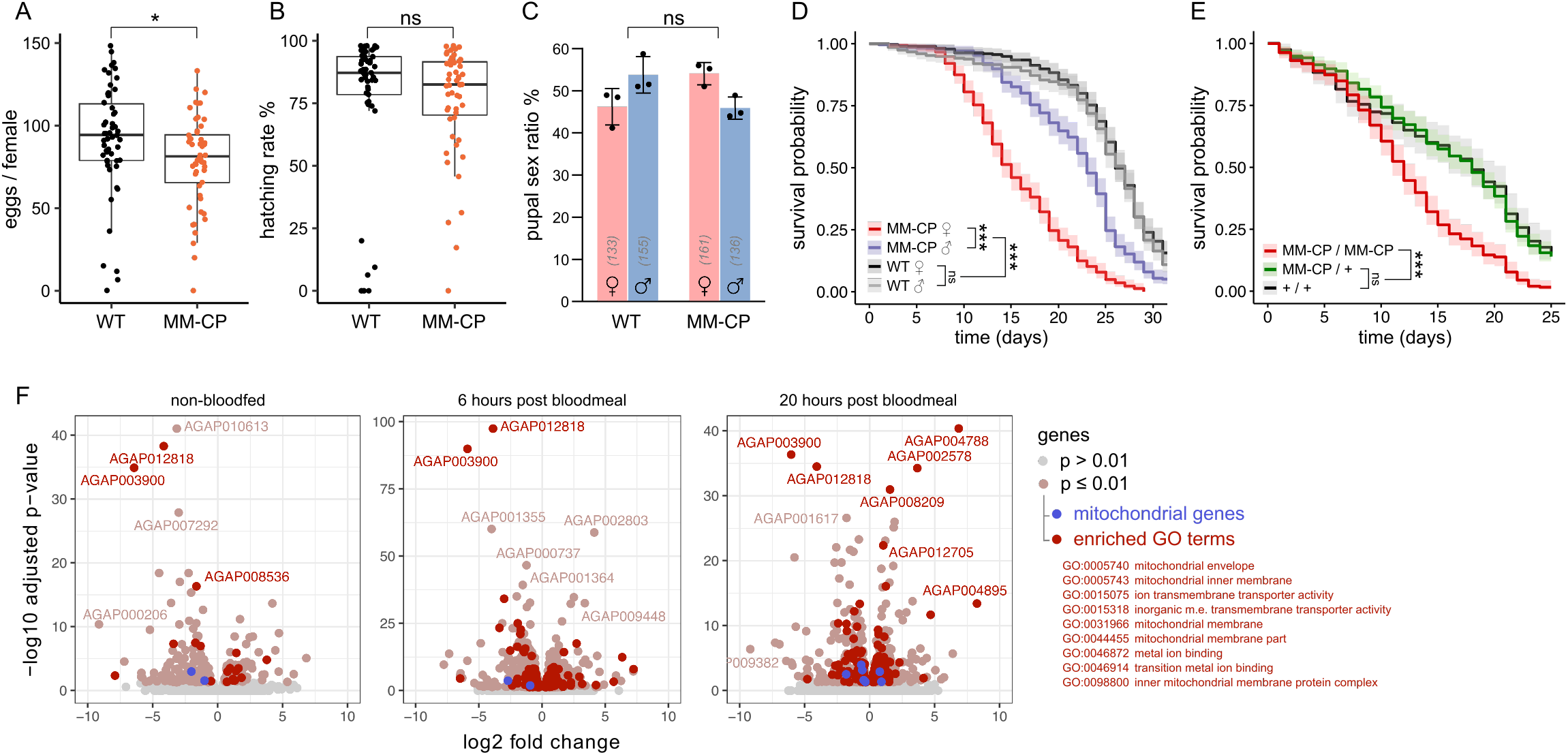
Life history traits and midgut transcriptome of MM-CP mosquitoes. **(A)** Fecundity of individual homozygous MM-CP females compared to the wildtype (WT) and corresponding **(B)** larval hatching rates obtained during the first gonotrophic cycle. Data from 3 pooled biological replicates are shown. Statistical significance was determined by a t-test assuming unequal variance. **(C)** Pupal sex ratios of MM-CP and WT strains analyzed using the χ^2^ test for equality. **(D)** Survival analysis of MM-CP and WT male and female mosquitoes maintained on sugar and (**E**) of F2 genotyped MM-CP female mosquitoes following backcrossing to the Ifakara strain, intercrossing of F1 mosquitoes and provision of bloodmeals. Statistical significance was determined with a Mantel-Cox log rank test. Data from three biological replicates are pooled and the mean and 95% confidence intervals are plotted. Significance codes *(p≤0.05), **(p≤0.01), ***(p≤0.001) and ns (not significant). **(F)** Volcano plots of an RNAseq experiment performed on midguts dissected before or 6 hours and 20 hours post bloodmeal. Differentially expressed genes between MM-CP and WT mosquitoes (p≤0.01) are indicated, and genes belonging to enriched GO groups are highlighted in red.

We performed transcriptomic analysis of dissected midguts prior to and 6 and 20 hours after blood feeding. We quantified the number of differentially expressed genes between MM-CP and control females and performed a gene ontology (GO) analysis to determine significantly enriched gene groups (**Table S1**). The results indicated that genes involved in mitochondrial function and located at the inner mitochondrial membrane were disproportionally affected in MM-CP females, particularly after the bloodmeal (**Fig. 3F–H**). Among most significant hits were genes encoding a member of the ubiquinone complex (AGAP003900), a mitochondrial H^+^ ATPase (AGAP012818), an ATP synthase subunit (AGAP004788) and a protein belonging to a family of calcium channels (AGAP002578), which control the rate of mitochondrial ATP production. These findings offered a hypothesis that could explain the dual phenotype regarding parasite development and adult female lifespan. *Plasmodium* development in the mosquito is critically dependent on mitochondrial function including active respiration (*45–48*). Magainin 2 and Melittin are known to trigger mitochondrial uncoupling and could, upon secretion into the midgut lumen, interfere with ATP synthesis targeting the parasite mitochondrion. This effect would become apparent as the parasite transforms into the energy-demanding oocyst stage that undergoes several rounds of endomitosis and vegetative growth. As AMPs are unlikely to be able to access the oocyst, the effect would wear off with time and indeed partly offset by supplemental bloodmeals. The AMPs, however, are likewise expected to affect the mosquito midgut mitochondria, impacting energy homeostasis and modulating lifespan. Whilst further experiments are needed to untangle these effects, the most significant knowledge gap, when it comes to transmission blocking, is to what degree any effects observed with lab strains of *P. falciparum* would be reproducible in infections with genetically diverse parasites isolated from patient blood. The MM-CP strain is an excellent candidate to attempt to answer this question as the transmission-blocking mechanism we describe here appears to act across *Plasmodium* species. MM-CP is incapable of autonomous gene drive, unless it mates with a mosquito source of Cas9, and can thus be evaluated in an endemic setting under standard mosquito confinement protocols.

Finally, we predicted how deployment of the MM-CP effector trait would modify malaria epidemiology using a mechanistic, agent-based model of *P. falciparum* transmission that includes vector life cycle and within-host parasite and immune dynamics. The model is based on the EMOD framework that has recently been updated to enable the simulation of gene drives (*49*). There remain knowledge gaps that preclude a direct translation of experimental entomological or molecular data into epidemiological parameters, for example linking the observed reduction in sporozoite DNA and its quantitative effect on onward transmission. For the phenotypic effects, we thus estimated likely parameter value ranges (a 30-70% increase in time until sporozoites are released and a 40-100% reduction in infectious sporozoites) that we considered to be within physiologically plausible limits supported by our *in vivo* experiments. As a final parameter for the model, we experimentally determined the rate of non-autonomous gene drive of the MM-CP allele by pairing it with a source of Cas9, which resulted in high levels of homing of the transgene in both males and females: 96.01% and 98.91%, respectively (**Fig. 4A**). MM-CP, as a non-autonomous effector, could be flexibly deployed in conjunction with non-driving, self-limiting or, as we assumed in our model, a fully autonomous Cas9 gene drive that is able to mobilize MM-CP **(Fig. S4)**. We determined the probability of elimination by the last year of simulation, defined as the number of simulations per parameter set that have zero prevalence in the last year divided by the total number of simulations. Incidence reduction was evaluated for the duration of the simulation following gene drive releases compared to control scenarios with no releases. In a low transmission setting (annual EIR ~15 infectious bites per person), most simulations within the space of likely parameter estimates resulted in the elimination of malaria transmission (**Fig. 4B** & **Fig. S5A**). As transmission intensity increased (annual EIR ~30), we detected a reduction in probability of elimination in the lower end of the parameter estimate range (**Fig. 4B** & **Fig. S5B**). In a high transmission scenario, a high reduction in sporozoite production in combination with a large delay was necessary to reliably trigger elimination (**Fig. 4B** & **Fig. S5C**). It should, however, be noted that significant reductions in clinical cases occurred even when elimination was not achieved. Therefore, in high transmission settings, even when not achieving elimination alone, MM-CP could open a window for elimination by strategically deploying other interventions that could act synergistically to drive transmission to zero.

**Figure 4.**
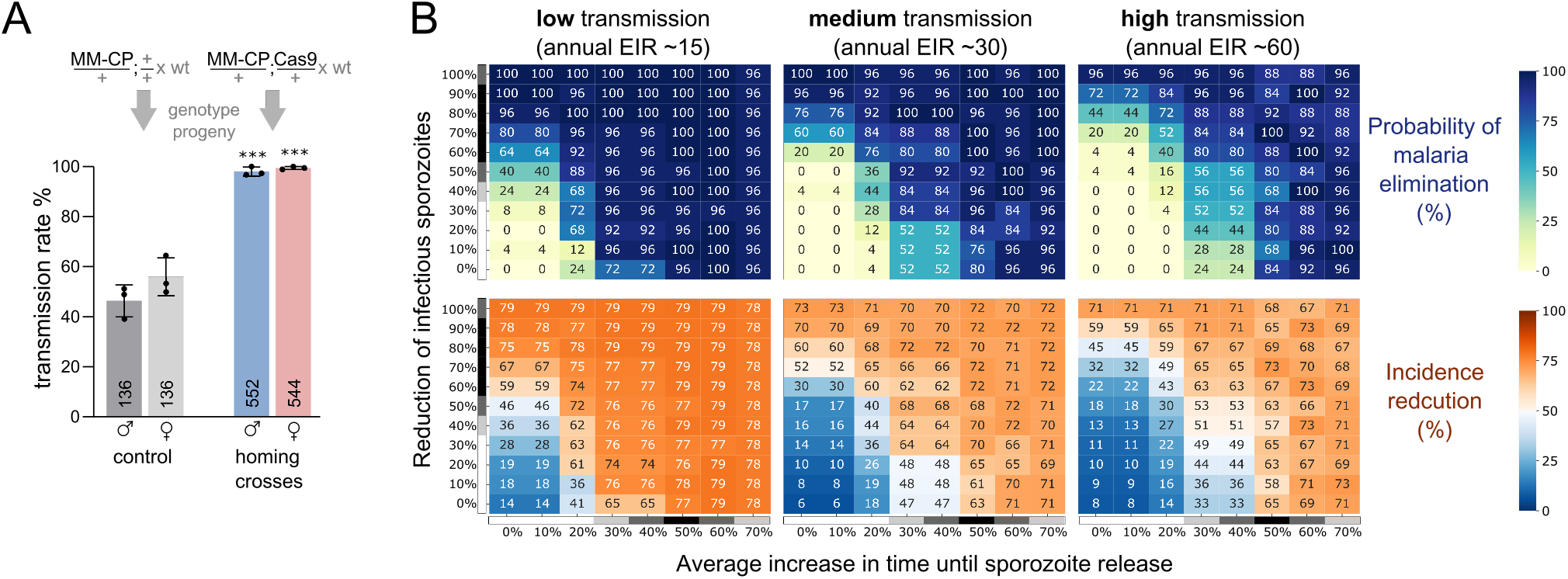
Gene drive and predicted epidemiological impact of strain MM-CP deployment. **(A)** Assessment of non-autonomous gene drive in the progeny of male and female hemizygous MM-CP mosquitoes in the presence or absence of a vasa-Cas9 driver crossed to the wildtype (wt). Larval offspring were subjected to multiplex PCR genotyping and the mean and standard error (SEM) from three biological replicates is plotted, and the total number n is indicated. Statistical significance was determined using a one-way ANOVA with Tukey’s correction. Significance codes *(p≤0.05), **(p≤0.01), ***(p≤0.001) and ns (not significant). **(B)** Heatmaps depicting elimination probabilities (top) and number of clinical cases reduced (bottom) at the end of 6 years following a single release of 1000 homozygous MM-CP mosquitoes that also carry a Cas9 integral gene drive. Three transmission scenarios with varying entomological inoculation rates (EIR) for *P. falciparum* as a measure of exposure to infectious mosquitoes were explored. Homozygous transgenic mosquitoes are released 6 months after the start of the simulation in highly seasonal transmission settings of varying intensities. The parameter range we explored for the reduction of the number of infectious sporozoites is represented on the major y-axis while the range for the average increase in time until sporozoites are released is represented on the major x-axis. Parameter likelihood estimates based on the experimental data are also indicated next to the values (grayscale).

## Acknowledgments

We thank Eric Marois for sharing the vasa-Cre and vasa-Cas9 lines. We thank Dan Bridenbecker for software support and Austin Burt for helpful suggestions on the manuscript.

## Materials and Methods

### Design and generation of constructs

Annotated DNA sequence files for the final transformation constructs pD-Gam1-MM and pD-MM-CP are provided in Supplementary file S1. Briefly, the 23 N-terminal amino acids of Cecropin 1 (Cec1, CecA, AGAP000693) and Cecropin 2 (Cec2, CecB, AGAP000692) served as secretion signals. Magainin 2, Melittin, T2A and P2A were codon usage optimized for *Anopheles gambiae*, and the intron located within the Melittin coding sequence was previously described (*42*) except for the SV40 terminator within the eGFP marker module for which we swapped in the Trypsin terminator (*50*). We first neutralized a BsaI site in the Ampicillin resistance cassette and gene synthesized (Genewiz) a fragment ranging from the Cecropin 1 secretion signal to the Trypsin terminator, including 18 bp overlaps with the vector backbone and eGFP for subsequent Gibson assembly. The marker-module and the U6 promoter was PCR amplified from pI-Scorpine (*42*) with primers 78-GFP-R and 167-U6-R. The fragment from the BsaI spacer to the P2A was synthesized (Genewiz), including 18 bp overlaps to the U6 promoter and the vector backbone. The vector backbone was PCR amplified from pAmpR_SDM with primers 168-BBmut-F and 169-BBmut-R, and finally the 4 fragments were joined via Gibson assembly to yield the intermediate plasmid pI-MM. The CP gRNA spacer (*42*) was inserted via the BsaI sites and the cassette was amplified with primers 172-CP-HA3-F-degen and 173-CP-HA5-R-Cec1 and fused with the CP homology arms and backbone amplified from pD-Sco-CP (*42*) with primers 170-Cec1-SS-F and 171-P2A-R to assemble the donor plasmid pD-MM-CP. A 5’ P2A was added to the cassette via Golden Gate cloning of the annealed oligos 182-P2Aanneal-F and 183-P2Aanneal-R into BglII digested plasmid pI-MM, and subsequently the Gambicin 1 gRNA spacer (5’-*TACAGAATGTTTCTTCTGAG*-3’) was inserted via BsaI. The gRNA sequence was chosen using Deskgen (Desktop Genetics, LTD) with an activity-score of 54 and an off-target score of 99. The effector cassette was amplified with primers 184-P2Adegen-F and 185-Mel-R and fused via Gibson assembly with the Gambicin homology arms amplified from G3 genomic gDNA and the backbone to generate the donor plasmid pD-Gam1-MM. For primers see Supplementary Table S2

### Transgenesis and establishment of markerless strains

#### Anopheles gambiae

G3 eggs were injected with the corresponding donor plasmids pD-MM-CP or pD-Gam1-MM and the Cas9 helper plasmid p155 (*4*). 25 F1 transgenics were obtained for MM^GFP^-CP and one female F1 transgenic for Gam1-MM^GFP^. MM^GFP^-CP was established from a founder cage with 9 females and F1 individuals were confirmed by Sanger-sequencing with primers EGFP-C-For, 117-CP-ctrl-R and 163-P3-probe-F, and Gam1-MM^GFP^ with primers EGFP-C-For and EGFP-N. Transgenics were outcrossed to G3 WT over three generations for Gam1-MM^GFP^ and two generations for MM^GFP^-CP before crossing to the vasa-Cre (*51*) strain in the KIL background. Larval offspring were screened for GFP and DsRed and siblings mated. The progeny was screened against GFP and DsRed, pupae were singled out and the exuviate collected for genotyping with primers 99-CP-locus-F and 100-CP-locus-R or 241-Gam-locus-F and 242-Gam-locus-R, respectively, to identify homozygotes. The markerless line MM-CP was established from 9 males and 11 females. For Gam1-MM, 3 cups with 1 female and 1 male and 6 cups with 2 males and 2 females were set-up and pooled after confirmation via Sanger-sequencing. A G3-KIL mixed colony was used as WT control for all experiments, unless otherwise stated. All experiments were performed with cow blood (First Link (UK) Ltd.), unless otherwise stated.

### RT-PCR and splicing analysis

MM-CP and the WT control were fed with human blood and midguts were dissected after 3h. For Gam1-MM this experiment was performed on unfed females. 30 guts were lysed in Trizol and homogenized with 2.8mm ceramic beads (CK28R, Precellys) for 30s at 6,800rpm in a Precellys 24 homogenizer (Bertin). RNA was extracted with the Direct-zol RNA Mini-prep kit (Zymo Research) including on-column DNase treatment and transcribed into cDNA with the qScript cDNA Synthesis Kit (Quantabio). RT-PCR was performed with Phire Tissue Direct PCR Master Mix kit (Thermo Scientific) using primers 429-Gambicin-F & 430-Gambicin-R, 270-qCP-F1 & 271-qCP-R3, 484-qMag-both-F & 485-qMel-both-R, as well as 447-S7-F & 448-S7-R for the S7 reference gene. To quantify splicing efficiency, PCRs were performed on above cDNAs with Q5 High-Fidelity DNA Polymerase (NEB) using 484-qMag-both-F as forward primer and 242-Gam-locus-R (365bp amplicon) or 246-qCP-R2 (309bp) as reverse primers, respectively. Annealing temperature, extension time and cycle number were set to 67°C, 5 seconds and 27 cycles, respectively. Amplicons were purified with QIAquick PCR Purification Kit (Qiagen) and submitted to Amplicon-EZ NGS (Genewiz) and the data (NAR accession PRJNA778891) analysed using Geneious Prime (Biomatters).

### Mosquito infection assays

Transgenic or control mosquitos were infected with mature *P. falciparum* NF54 gametocyte cultures (2-6% gametocytaemia) as described previously using the streamlined Standard Membrane Feeding Assay (*29*) or with *P. berghei* ANKA 2.34 that constitutively expresses GFP by direct feeding on infected mice. Engorged mosquitoes were provided 10% sucrose and maintained at 27 °C for *P. falciparum* infections and 21 °C/RH 75% for *P. berghei* infections until dissections were performed. Supplemental bloodmeals on human blood was provided via membrane feeding. For infections of mosquitoes with mature *Plasmodium falciparum* mosquitoes were starved without sugar for 48 hours after the infective or supplemental bloodmeal to eliminate unfed individuals.

### Analysis of parasite infection intensity and prevalence

We dissected midguts at the indicated days and microscopically examined them for the presence of oocysts after staining with 0.1% mercurochrome. We measured the diameter of oocyst using ImageJ (*52*). For measuring the prevalence and intensity of sporozoites, the head and thorax as well as the midgut were dissected for each female. The gDNA was extracted separately from head/thorax samples and the corresponding midgut samples with the DNeasy 96 Blood and Tissue Kit (Qiagen) and was used for qPCR 20μl reactions using the Qiagen Quantinova SYBR Green PCR kit to quantify the *P. falciparum* Cytochrome B gene fragment using primers and methods described previously (*43, 53*). Standard curves for the target gene and the *A. gambiae* S7 reference gene were calculated after serial dilution of nucleic acid templates. Ct-values were converted using their respective standard curves, and the target gene Ct value was normalized to the reference gene (*A. gambiae* S7 ribosomal gene).

### Generation of backcross populations and genotyping

50 homozygous MM-CP males and females were crossed to 50 *Anopheles gambiae s.s*. Ifakara strain females and males respectively. F1 siblings were mass mated in a single cage to obtain F2 progeny. F2 females were used for infection experiments or survival assays as described. For genotyping of individual mosquitoes, we used we used multiplex genomic PCR with primers CP-multi-F, CP-multi-R and Mag-R which results in an 356bp amplicon for the MM-CP transgene and 670bp for the unmodified CP locus.

### Fitness and survival assays

For each replicate 20 females were individually transferred to cups one day after being offered an uninfected bloodmeal. Spermatheca were then dissected from females that failed to oviposit eggs to exclude unfertilized individuals. Eggs and larvae were counted on day 7 after blood-feed. To determine the pupal sex ratio and pupation time, 100 L1 larvae per tray were reared to the pupal stage where pupae were being collected and sexed once a day. Three biological replicates were performed, and the data were analysed via the Chi-square test for deviations from the expected sex ratio of 50%. The average pupation time in days was calculated and tested for statistical significance with the Mann-Whitney test. For the survival analysis including both sexes, a total of 274 WT and 276 MM-CPmale and 275 WT and 304 MM-CP female pupae were placed in 6 separate cages with bottles containing 10% filtered fructose solution and accumulated dead mosquitoes were counted daily. Survival was monitored daily for 44 days on 3 independent replicates. For the survival analysis of backcrossed females F2 mosquitoes were placed in W24.5 x D24.5 x H24.5 cm cages as pupae. Mosquitoes were offered a 10% sugar solution and they were also offered a blood meal and allowed to deposit eggs 72 hrs after every bloodmeal. Dead mosquitoes were collected every 24 hrs from the cage and preserved to be genotyped. Survival was monitored daily for 25 days on two independent replicates.

### RNAseq analysis

Females were fed with human blood and 15 guts were dissected into Trizol after 6 hours, 20 hours and from unfed females. After homogenization with 2.8mm ceramic beads (CK28R, Precellys), RNA was extracted with the Direct-zol RNA Mini-prep kit (Zymo Research) including on-column DNase treatment. Four biological replicates per condition were subjected to RNA-seq. Libraries were prepared with the NEB Next® Ultra™ RNA Library Prep Kit and sequenced on a NovaSeq 6000 Illumina platform (instrument HWI-ST1276) generating 150bp paired-end reads (NAR accession tbc). Replicate 1 for MM-CP without bloodmeal (MMCP_N_1) was identified as outlier with squared Pearson correlation coefficients with the other three biological replicates below 0.84, and hence removed from further analysis. Sequencing reads were mapped to the Anopheles gambiae PEST genome (AgamP4.13, GCA_000005575.2 supplementred with the MM-CP construct reference) using HISAT2 software v2.0.5 (*54*) (with parameters --dta --phred33). Differential expression was assessed with DESeq2 v1.20.0. and GO enrichment analysis was performed using TopGO (*55*) with a pruning factor of 50 using a p-value cutoff of p=0.01.

### Assessment of non-autonomous gene drive

At least 60 homozygous MM-CP or wild type females were crossed to males of the vasa-Cas9 strain. F1 progeny were screened for the presence of the 3xP3-YFP marker and transhemizygotes were then sexed and crossed to wild types. Genomic DNA was isolated from the progeny at the L2-L3 larval stage according to the protocol of the Phire Tissue Direct PCR Kit (Thermo Scientific). Multiplex PCR was performed with primers 260-q-Mag-Mel-R, 531-CP-multi-R and 532-CP-multi-F, yielding a 356bp band if the construct is present and a WT band of 670bp as control. Two 96-well-plates per parent (paternal or maternal transhemizygotes) and replicate were analysed and four negative controls were included on each plate. From the control-crosses, 46 offspring per parent were analysed for each replicate. The homing rate was calculated as (n*0.5 – E_neg_) / (n*0.5) *100, with E_neg_ being the individuals negative for the effector and n being the total number of samples successful analysed by PCR.

### Transmission modelling using EMOD

Simulations were performed using EMOD v2.20 (*56*), a mechanistic, agent-based model of *Plasmodium falciparum* malaria transmission that include vector life cycle dynamics and within host parasite and immune dynamics. Seasonality of rainfall and temperature as well as vector species were kept the same across transmission settings, but vector density was varied to match desired transmission intensity. *A. gambiae*, the only vector considered, was assumed as being 95% endophilic and 65% anthropophilic. Each simulation contained 1000 representative people with birth and death rates appropriate to the demography without considering importation of malaria. We include baseline health seeking for symptomatic cases as an intervention where human agents can seek treatment with artemether-lumefantrine (AL) 80% of the time within 2 days of severe symptom onset and 50% of the time within 3 days of the onset of a clinical but non-severe case. Mosquitoes within EMOD contain simulated genomes that can model up to 10 genes with 8 alleles per gene with phenotypic traits that map onto different genotypes (*49*). Here we modelled an integral gene drive system (*57*) aimed at population replacement with an effector that results in delayed sporozoite production in infected mosquitoes as well as an overall reduction in the number of sporozoites produced by infected mosquitoes. The model was further parameterized using the experimentally determined measures of fitness, lifespan and homing. 1000 male IGD mosquitoes homozygous for the autonomous drive (Cas9) and non-autonomous effector (MM-CP) were released just before the wet season begins to pick up in transmission intensity (Table S2). Apart from the sex chromosomes, two loci representing the effector and driver were modelled (*57*), with each locus having four possible alleles (wild type, resistant, effector or nuclease and loss of gene function for the effector or driver locus). To evaluate the performance of these drives in a range of transmission settings, we vary transmission intensity via annual entomological rates (EIR) ranging from 15 infectious bites per person to 60 infectious bites per person. We also vary the final phenotypic effect of expressing the effector gene that leads to delayed and reduced sporozoite formation. We evaluate average increases in time until sporozoite formation ranging from no increase in time compared to a wild-type mosquito up to 70% increase in sporozoite formation time. As for the reduced sporozoite effect, we evaluated to full range of possible effects compared to a wild-type mosquito. Mosquitoes carrying the drive are released 6 months into the simulation, and simulations are run for a total of 6 years. The outputs represent the mean of 25 stochastic realizations per parameter set.

---

**Fig. S1.**
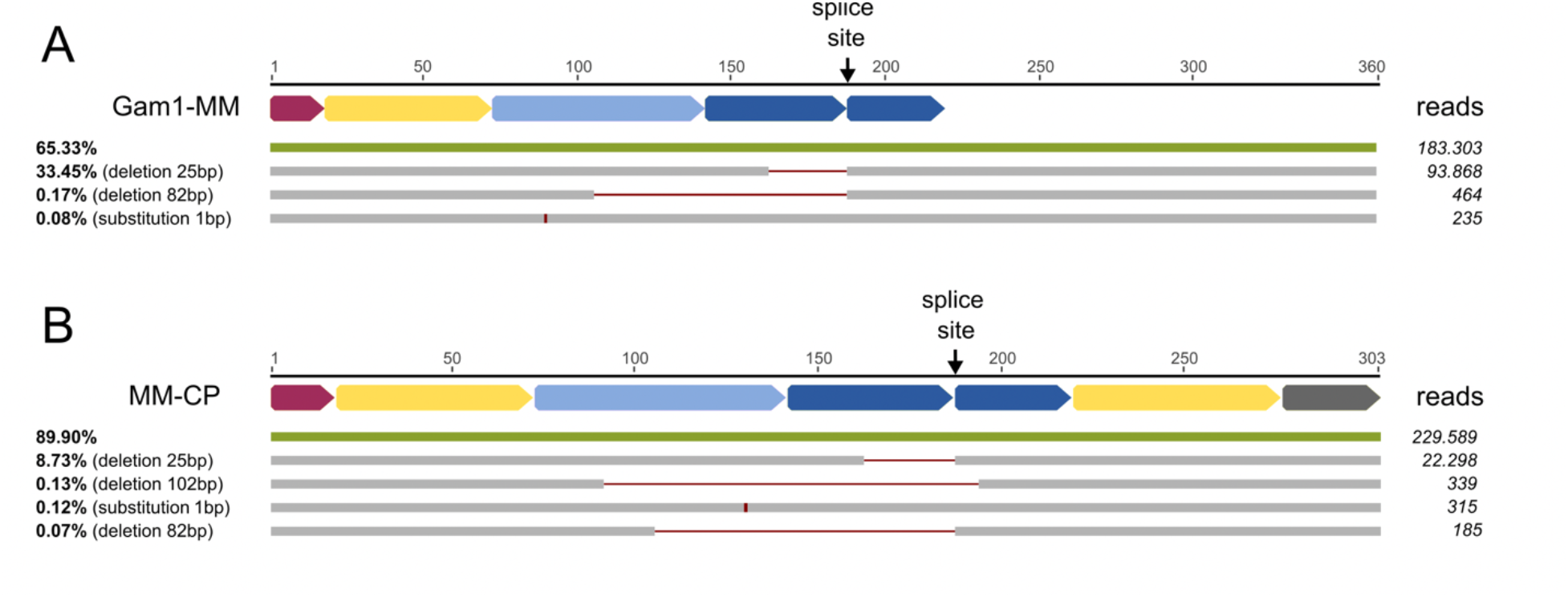
Alignment of sequenced cDNA amplicons to reference sequences representing the expected splicing outcomes for Gam1-MM and MM-CP. The splice site is indicated by the black arrow, the relative distribution of predicted variants (grey) representing at least 0.07% of all reads is shown on the left and the corresponding number of reads on the right.

**Fig. S2.**
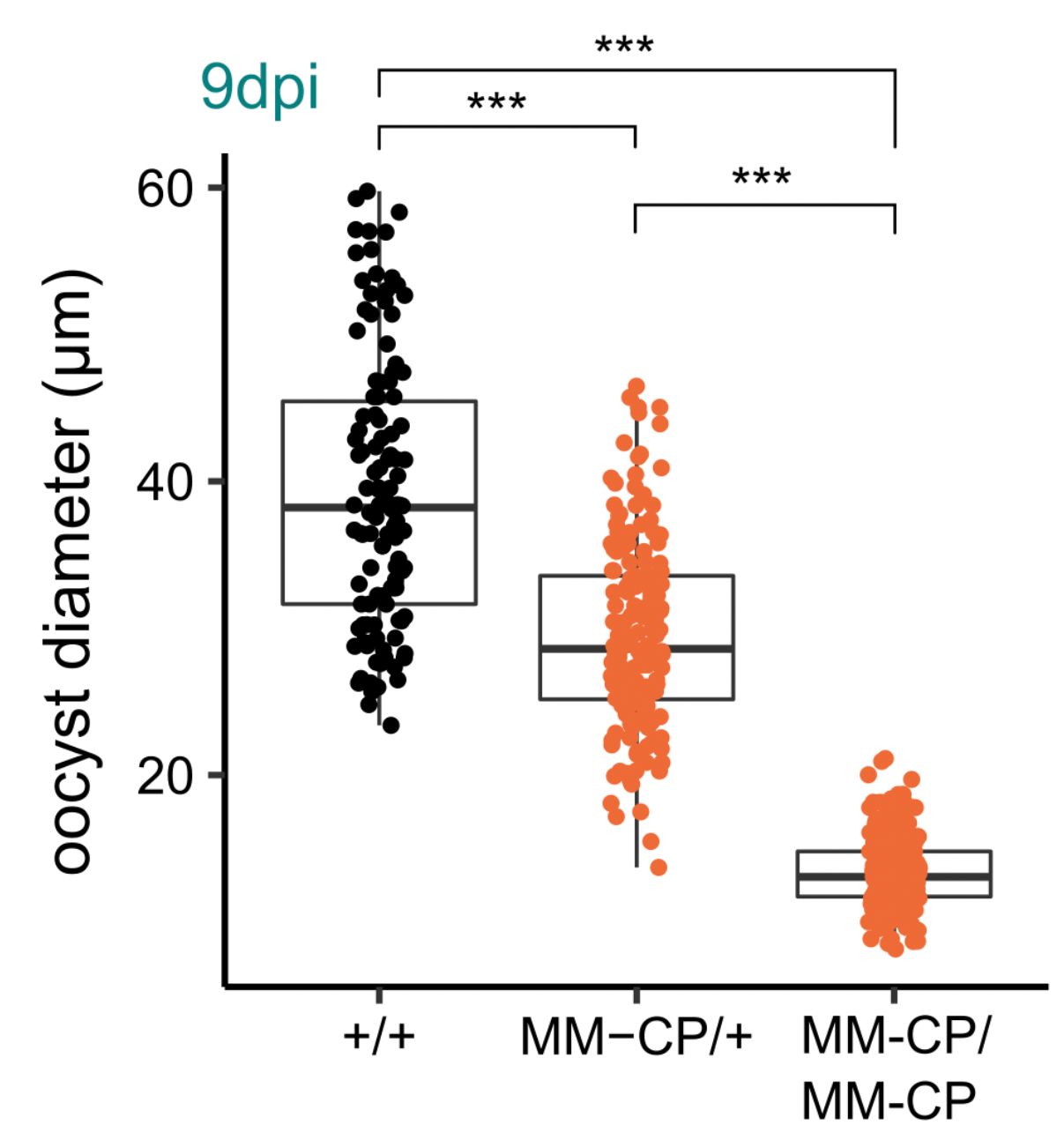
Quantification of *P. falciparum* oocyst diameter in F2 individuals following a backcross of strain MM-CP to the Ifakara strain and an F1 sibling intercross from 3 pooled biological replicates. Non-transgenic (+/+), hemizygous (MM-CP/+) and homozygous (MM-CP/MM-CP) individuals were identified by individual PCR genotyping after oocyst size had been determined. Statistical analysis was performed by a t-test assuming unequal variance.

**Fig. S3.**
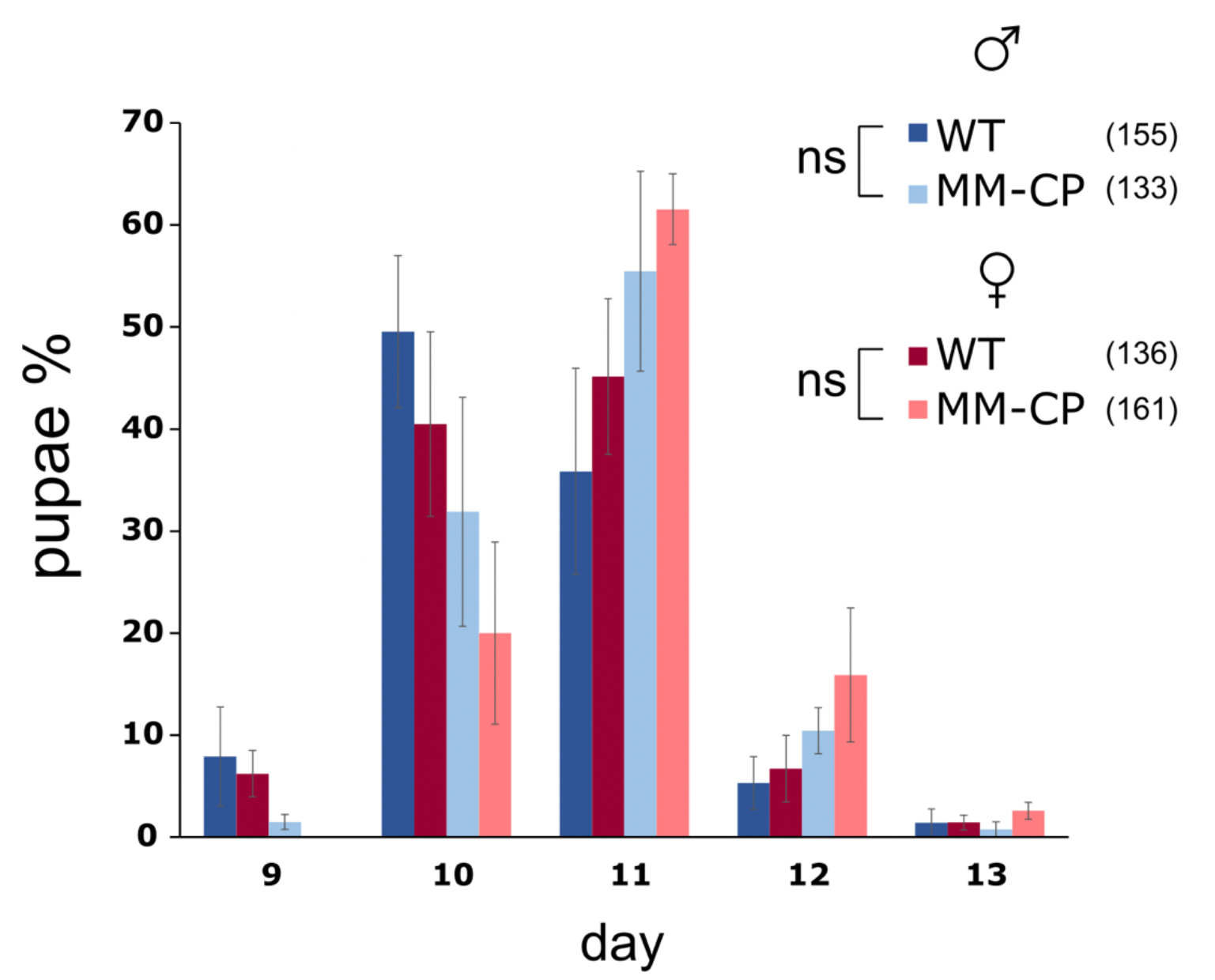
Analysis of the time of pupation of MM-CP and wild-type mosquitoes as the percentage of pupae emerging each day. Statistical analysis of average pupation times in males and females was calculated using the Mann-Whitney test.

**Fig. S4.**
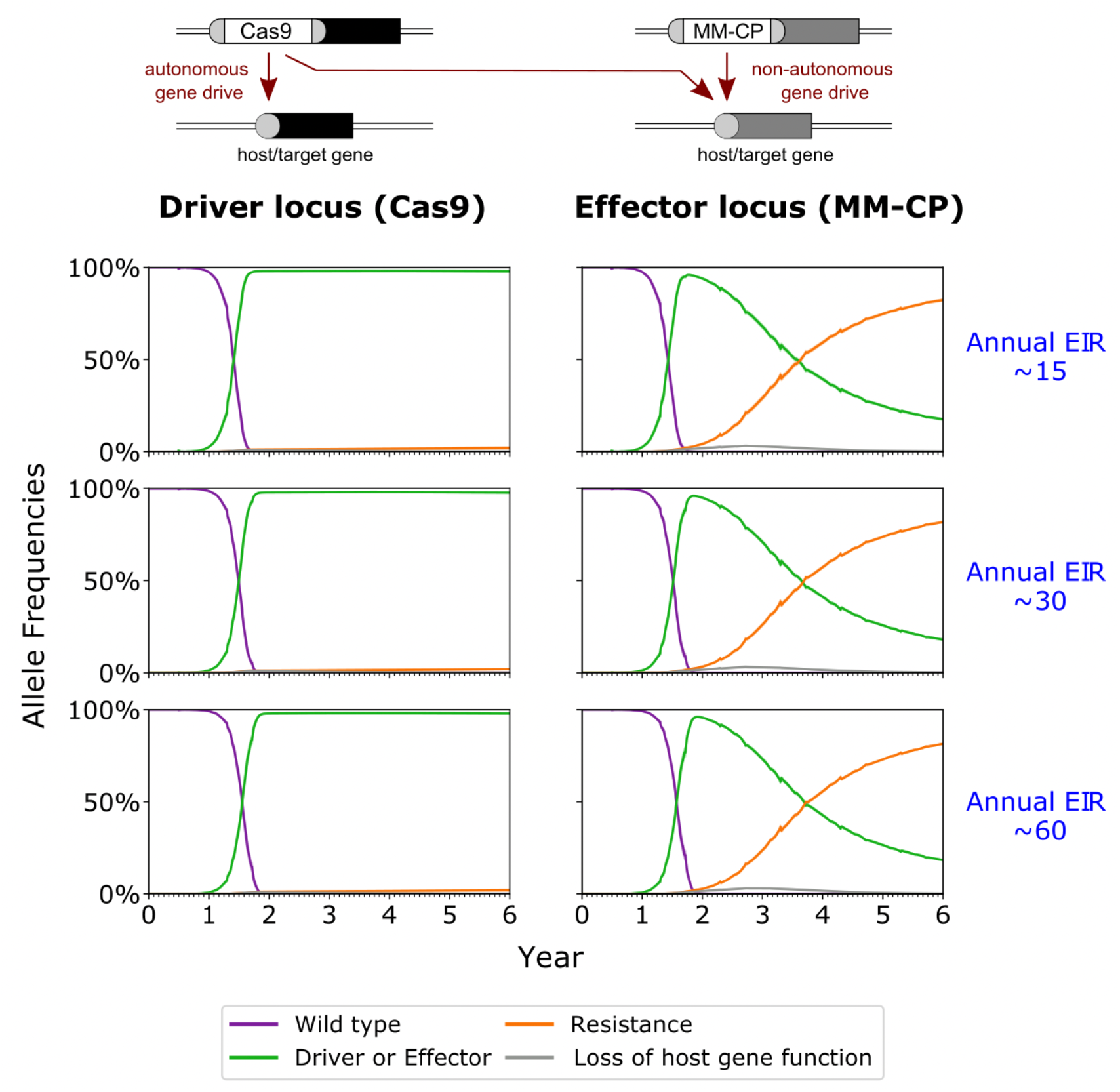
The mean time course of allele frequencies after 1000 gene drives mosquitoes homozygous for the driver and effector locus are released 6 months into a 6-year simulation. Each row represents a different transmission intensity. The left column depicts allele frequency at the driver locus and the right column represents allele frequencies at the effector locus. Shaded regions represent one standard deviation on either side of the mean allele frequency as calculated from 25 stochastic realizations of each scenario.

**Fig. S5.**
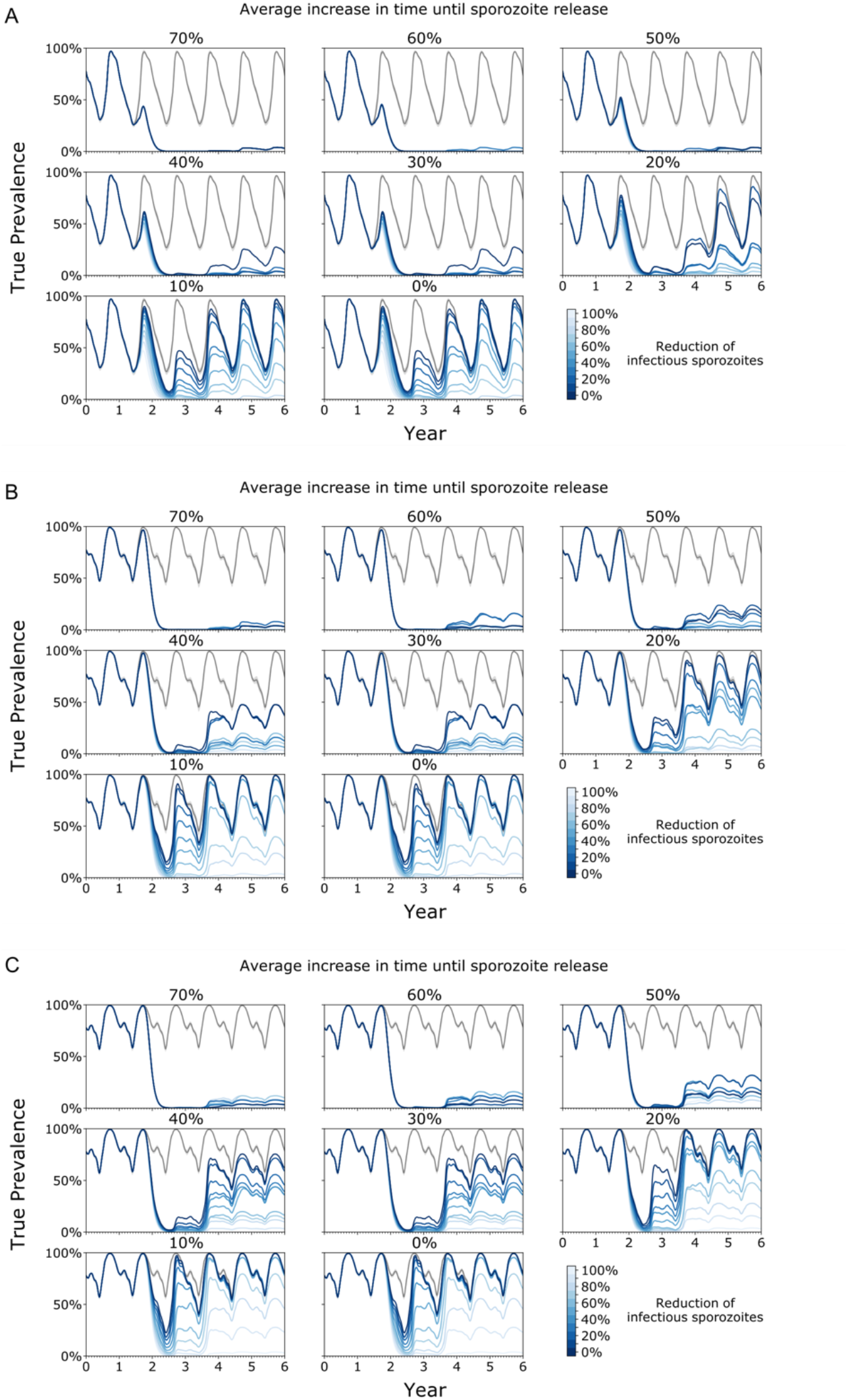
The mean time course of true prevalence after 1000 gene drives mosquitoes homozygous for the driver and effector locus are released 6 months into a 6-year simulation. Annual EIR of around 15 (A), 30 (B) and 60 (C) infectious bites per person in an unmitigated scenario. Each panel represents the average increase in time until sporozoites are released while the blue shaded traces each represent a different average reduction in infectious sporozoites. The grey trace represents the unmitigated baseline scenario.

**Table S1.** Enriched GO terms

| GO.ID | Term | Condition | gene counts |  |  | p value |  |
| --- | --- | --- | --- | --- | --- | --- | --- |
|  |  |  | Annotated | Significant | Expected | elimKS | classicFisher |
| GO:0046914 | transition metal ion binding | 6 hours | 95 | 72 | 59.41 | 0.004 | 0.0031 |
| GO:0046872 | metal ion binding | 6 hours | 201 | 141 | 125.71 | 0.106 | 0.0086 |
| GO:0015318 | inorganic m.e. transmembrane transporter activity | 20 hours | 52 | 38 | 27.52 | 0.00019 | 0.0018 |
| GO:0015075 | ion transmembrane transporter activity | 20 hours | 59 | 41 | 31.23 | 0.00046 | 0.0057 |
| GO:0098800 | inner mitochondrial membrane protein complex | 20 hours | 56 | 39 | 29.68 | 0.0021 | 0.0066 |
| GO:0031966 | mitochondrial membrane | 20 hours | 78 | 52 | 41.34 | 0.4586 | 0.0073 |
| GO:0005740 | mitochondrial envelope | 20 hours | 80 | 53 | 42.4 | 0.4701 | 0.0082 |
| GO:0044455 | mitochondrial membrane part | 20 hours | 65 | 44 | 34.45 | 0.384 | 0.009 |
| GO:0005743 | mitochondrial inner membrane | 20 hours | 72 | 48 | 38.16 | 0.435 | 0.0099 |

**Table S2.** Table of EMOD modelling parameters.

| *Fitness & phenotypic parameters* |  |  |  |  |
| --- | --- | --- | --- | --- |
| Locus | Allele combination | Daily mortality (%) | Fecundity (%) | Applied to sex |
| driver | WT , D |  | -2.5 | M,F |
| driver | WT , LGF |  | -10 | M,F |
| driver | D , D |  | -5 | M,F |
| driver | D , R |  | -2.5 | M,F |
| driver | D , LGF |  | -12.5 | M,F |
| driver | R , LGF |  | -10 | M,F |
| driver | LGF , LGF |  | -100 | M,F |
| effector | WT , LGF | +10 |  | M,F |
| effector | E , E | +30 | -14 | F |
| effector | E , LGF | +10 |  | M,F |
| effector | R , LGF | +10 |  | M,F |
| effector | LGF , LGF | +100 |  | M,F |
| *Gene drive parameters* |  |  |  |  |
| Locus | Allele combination | Outcome | Probability | Allele generated |
| driver | D , <u>WT</u> | Gene Drive (autonomous) | 0.97 | D |
|  |  | Unmodified | 0.1 | WT |
|  |  | NHEJ to R2 | 0.1 | LGF |
|  |  | NHEJ to R1 | 0.1 | R |
| effector | E , <u>WT</u> (+D ) | Gene Drive (non-autonomous) | 0.97 | E |
|  |  | Unmodified | 0.1 | WT |
|  |  | NHEJ to R2 | 0.1 | LGF |
|  |  | NHEJ to R1 | 0.1 | R |

**Supplementary file S1. (separate file)**

Annotated DNA sequences of transformation vectors pD-MM-CP and pD-Gam1-MM.

**Supplementary file S2. (separate file)**

DNA oligonucleotide database.

